# Multi-Omics Analysis of Wound Microbiome and *Staphylococcus aureus* in Pressure Ulcer

**DOI:** 10.1101/2025.02.08.637207

**Authors:** Changyuan Guo, Guilai Jiang, Yuanyuan Li, Jun Zhao, Jilin Li, Shengkai Li, Zhemin Zhou, Erfan Xie, Heng Li

**Affiliations:** Suzhou BenQ Medical Center, Suzhou, Jiangsu Province, China; Department of Microbiology, School of Basic Medical Sciences, Suzhou Medical College, Soochow University, Suzhou 215123, China; Key Laboratory of Alkene-Carbon Fibres-Based Technology & Application for Detection of Major Infectious Diseases, MOE Key Laboratory of Geriatric Diseases and Immunology, Cancer Institute, Suzhou Medical College, Soochow University, Suzhou 215123, China; Suzhou Key Laboratory of Pathogen Bioscience and Anti-infective Medicine, Jiangsu Province Engineering Research Center of Precision Diagnostics and Therapeutics Development, Soochow University, Suzhou 215123, China; Suzhou Key Laboratory of Smart Nursing and Health Care for the Elderly, Soochow University, Suzhou 215123, China

**Keywords:** pressure ulcers, *Staphylococcus aureus*, multi-omics, biofilm, metagenomic sequencing, antimicrobial resistance

## Abstract

This study employs a multi-omics approach to investigate the wound microbiome and *Staphylococcus aureus* in pressure ulcers. Metagenomic sequencing and associated technologies were utilized to examine bacterial populations, identify dominant species, and study biofilm formation in *S. aureus* through metabolomic and proteomic characteristics. The results revealed a significant reduction in microbial diversity in pressure ulcer samples compared to controls, with *Staphylococcus, Corynebacterium*, and *Klebsiella* being the most prevalent genera. Functional prediction analysis indicated differences in pathogenic invasiveness and biofilm formation factors among the groups. The presence of antimicrobial resistance genes was higher in the pressure ulcer group, particularly in *Staphylococcus* spp. strains. Whole-genome sequencing of 29 *S. aureus* isolates identified various clonal complexes and *spa* types, with the majority possessing genes conferring resistance to β-lactam antimicrobials and virulence factors. Histopathological examination and fluorescent in situ hybridization confirmed the presence of *S. aureus* in biofilm structures. In vitro biofilm formation tests and metabolomic and proteomic analyses provided insights into the interactions between *S. aureus* strains and their biofilm formation, revealing enriched pathways related to metabolic processes and membrane composition. This research offers a scientific foundation for understanding the colonization patterns of *S. aureus* biofilms in pressure ulcer wounds.

## Introduction

Among the most formidable threats to worldwide health is the delayed healing, especially pressure ulcers, impacting approximately 100 million individuals each year [1]. A variety of factors affect the recovery process including shifts in inflammatory reactions and skin microbiome. The interactions among bacteria within biofilms are crucial for understanding the persistence of infections, which pose a significant barrier to wound healing and hold substantial clinical relevance, especially in the context of an aging population[2-4].

By the late 19th century, there was a concerted global effort by scientists to isolate and characterize bacteria from biofilms found in wounds [5]. The chronic wounds, including pressure ulcers, tend to form biofilms and significantly reduced microbial diversity. A study by James et al. underscores this difference by comparing 50 chronic wound samples with 16 acute wound samples, revealing that an alarming 60% of the chronic wounds harbored biofilms [6-8]. In the context of Chinese patients, previous studies focused on isolating and cultivating bacteria from wound samples. Among the more than 60 cultivable bacteria identified, conditional pathogens were identified in relation to pressure ulcers including *Staphylococcus aureus, Escherichia coli, Pseudomonas aeruginosa*, and *Proteus mirabilis* [9,10].

*Staphylococcus aureus* is a significant concern in chronic wound infections, especially among individuals with pressure ulcers [11,12]. A study conducted by Luuk et al. examined the skin microbiota of patients with pressure ulcers and found that *S. aureus* and unclassified *Enterococci* were the most prevalent in these wounds. This research was pioneering in uncovering changes in the skin microbiome associated with pressure ulcers [13]. Furthermore, Wolcott et al. employed 16S rRNA gene sequencing and found significant quantities of *S. aureus* and *P. aeruginosa* in pressure ulcers [14]. However, what about the functional genes associated with pressure ulcers, including those responsible for virulence and antimicrobial resistance? How do the proteomic and metabolic profiles of *S. aureus* change during biofilm formation? The answers to these questions are yet to be determined.

This research centers on pressure ulcers, utilizing metagenomic sequencing and associated technologies to examine the bacterial populations within the wounds, pinpoint the dominant species, and perform biofilm formation studies on *S. aureus* by employing the metabolomic and proteomic characteristics. This approach provided a scientific investigation into the microenvironment of pressure ulcer wounds.

## Methods

### Ethics approval

The experiment was strictly conducted according to the Guide for the Care and Use from the Research Ethics Committee of Soochow University (20210220). All procedures involving human participants were performed in accordance with ethical standards. Patients were given informed consent in the study.

### Medical center

This study was conducted at Suzhou BenQ Medical Center. A tertiary hospital was established at this medical facility in 2013. The center has a construction area of 160,000 square meters with a total investment of 1,500 beds. In 2020, the hospital outpatient visits exceeded 500,000 cases, annual hospitalizations surpassed 25,000 cases, and the annual number of surgical procedures reached 10,000 cases. The facility is responsible for the diagnosis, treatment, and care of patients in Suzhou, with a particular focus on emergency trauma, critical care medicine, and health management as its key areas of expertise.

### Sample collection

A total of 50 samples were collected from the medical center between March to May 2023, comprising individuals with swabs of both pressure ulcers area and healthy skin area. To protect patient privacy, we only recorded age and pressure ulcer grade. Prior to sampling, the area was meticulously cleaned, followed by using a cotton swab to gently scrape the surface and transfer into a 2mL centrifuge tube. In cases where surgical excision was necessary, the excised tissue along with the surrounding area was placed into a centrifuge tube and swiftly sent to the laboratory for analysis.

### Total DNA extraction

After being removed from the sampler, the stool was diluted 5x with molecular water and homogenized with a vortex. The stool suspension (250 μl) was used for DNA extraction. The genomic DNA was purified and isolated with MagaBio Soil/Feces Genomic DNA Purification Kit (Hangzhou Bioer Technology, Hangzhou, China). The extracted DNA was quantified at an A260/A280 nm ratio with the NanoDrop ND-1000 (Thermo Fisher SCIENTIFIC, USA). Then extraction from each sample was diluted to approximately 5□ng/μl and stored at −20 °C for 16S rRNA sequencing.

### 16S rRNA amplicon sequencing

The 12-bp barcoded primers synthesized by Invitrogen (Invitrogen, Carlsbad, CA, USA) were used to amplify the bacterial 16S rRNAV3-V4 fragments. The PCR mixture contained 25 μl reactions Taq (Takara Biotechnology, Dalian, China), 1 μl of each primer (10 mM), and 3 μl DNA (20 ng/μl) template (final volume: 50 μl). The PCR protocol was as follows: 94 °C for 30 s followed by 30 cycles of denaturation at 94 °C for 30 s, annealing at 52 °C for 30 s, and extension at 72 °C for 30 s; followed by a final extension at 72 °C for 10 min. The PCR products were subjected to 1% agarose gel electrophoresis and then sequenced on an Illumina Miseq (PE 300) in the MAGIGENE Genomic Institute. The PCR products were mixed according to the Instructions of the GeneTools Analysis Software (Version 4.03.05.0, SynGene).

### Metagenomic next-generation sequencing (mNGS)

The extracted DNA that passed the quality inspection were randomly fragmented into approximately 350bp fragments using ultrasonic disruptor. The entire library preparation process includes steps such as end repair and amplification. After library construction, initial quantification is performed using Qubit 2.0, and the library is diluted to 2ng/uL. Subsequently, the insert size of the library is assessed using the Agilent 2100 system, ensuring the quality of the library. The qualified library will undergo sequencing using Illumina PE150 technology generate raw data for subsequent data analysis.

### *Staphylococcus aureus* isolation

The diagnosis was conducted by trained clinical staff from the Department of Infectious Diseases at the medical center, according to the guidelines for the treatment of MRSA infection. Quality control strains were used to ensure that contamination from various bacteria was minimized, including *Staphylococcus aureus* (ATCC29213), *Streptococcus pneumoniae* (ATCC9619), *Escherichia coli* (ATCC25922), and *Pseudomonas aeruginosa* (ATCC27853). The obtained S. aureus strains were confirmed by MALDI-TOF MS (BioMérieux, Craponne, France).

### Whole genome sequencing

Isolates were cultured for 24 h at 37°C in Tryptone Soya Broth (AOBOX 02-049, Beijing, China), and genomic DNA was extracted and purified using a HiPure Bacterial DNA Kit (D3146, Meiji Biotechnology Co., Ltd., Guangzhou, China). The extracted DNA was tested for quality using a Nanodrop ND-1000 spectrophotometer (Nanodrop Technologies, Wilmington, DE, USA) and a 1.0% (w/v) agarose gel. Purified DNA was whole genome sequenced via Illumina’s NextSeq 500 on the platform of Honsunbio company (Shanghai, China). The quality was assessed using QUAST v2.3 [15] and assembled by EToKi v1.0.

### Bioinformatics analysis

Quality control, assembly and subsequent phylogenetic analysis of sequencing data were performed using the program available in github (https://github.com/zheminzhou/EToKi), and the evolutionary tree was annotated and visualized using iTOL (https://itol.embl.de/). Based on VFDB, CARD database, Functional genes were identified based on VFDB and CARD database. Chao1, Shannon, Simpson, and ACE indices were computed using Qiime software (Version 1.9.1) [16], and Unifrac distances were calculated and UPGMA sample clustering trees were constructed. The results of PCA, PCoA, and Wilcoxon rank-sum test were visualized using R software (Version 4.1.2). PCA and PCoA analyses were performed using the stats and ggplot2 packages of the R software, and the Wilcoxon rank-sum test was performed using the ggplot2 package. Differences in microbial community composition between pressure ulcer samples and controls were compared using STAMP software (Version 2.1.3) [17]. Using PICRUSt2 [18], a bioinformatics software package for macrogenomic function prediction based on Marker genes, here we utilized PICRUSt2 and Bugbase tool [19] to perform KEGG database-based function prediction based on the sequencing data of the pre-existing 16S amplicons. Based on the functional annotations and abundance information of the samples in the database, the top 35 functional entries in terms of abundance and their abundance information in each sample were selected to draw heat maps and clustered at the level of functional differences. The sequences from mNGS were used for antimicrobial resistant gene prediction. The ARGs were predicted by ARGpore2 [20].

### Histopathological examination

All sections of pathological tissues were provided by the department of pathology. The project members only participated in the staining process. Simply, the tissues were fixed in 2.5% (w/v) glutaraldehyde-polyoxymethylene solution for 48□h. The fixed tissues were routinely processed, embedded in paraffin, sectioned (4□μm thickness), and stained with Hematoxylin and Eosin stain, Periodic Acid-Schiff stain, Gram staining, Masson staining, and Sirius-red staining.

### Fluorescent in situ hybridization

Targeting conserved regions of 16S rRNA and *S. aureus agrBDCA* operon, the 5′-end fluorescent motif-labeled probes were designed using the probe design function of the ARB program (www.ARB-home.de), and the colony probes were customized on Sangon.com. Configure hybridization solution [900 mM NaCl, 20 mM Tris, pH 7.5, 0.01% SDS, 20% formamide] at a final concentration of 2 nM for each probe. slides containing the biofilm were shaken and washed 3 times in PBS on a decolorizing shaker for 5 min each time. drops of hybridization solution were placed on the sample sections and hybridized overnight in a humid box at 37°C. Wash buffer 215 mM NaCl, 20 mM Tris, pH 7.5, 5 mM EDTA was configured. The slides are then washed with Wash Buffer at 37°C for 15 minutes, with additional formamide washes if there are more non-specific hybrids. Pour off the prehybridization solution, add a drop of probe 2 hybridization solution, and hybridize at 37°C overnight. The slices were incubated with DAPI staining solution for 8 min, rinsed, and sealed with a drop of fluorescence quenching sealer. Sections were viewed under a Nikon orthogonal fluorescence microscope and images were acquired. The images were reconstructed using ZEN software (version 8.1.0.484) based on the structured illumination algorithm with noise filter settings ranging from -6 to -8. The images were then analyzed using the COMSTAT plug-in in Image J. The images were then analyzed using the COMSTAT plug-in in Image J. The images were then analyzed using the COMSTAT plug-in in Image J.

### In vitro biofilm assessment

Inoculated 50ul bacterial suspension into a 96-well plate, added 50ul TSB medium, covered the plate, and incubated at a constant temperature of 37°C. After 72 hours of cultivation, discarded the contents of the 96-well plate, rinsed three times with 250 uL of sterile phosphate buffer (pH 7.3) to remove the culture medium and planktonic bacteria. Added 250 uL of methanol to each well, covered, let it stand for 15 minutes, aspirated the methanol, and air-dried at 30°C to fix the tightly adhered bacteria. In each well, added 250 uL of crystal violet (1% crystal violet solution), let it stand for 20 minutes, discarded the staining solution, and rinsed with sterile phosphate buffer (pH 7.3) until colorless; then air-dried. Added 250 uL of 95% ethanol to each stained well for destaining, used an empty well with 250 uL of TSB medium as a blank for zero adjustment, and measured the OD value at a wavelength of 595 nm. Performed triplicate measurements for each well and calculated the average.

### In vitro metabolomic analysis of biofilms

Sample preparation involved thawing samples on ice, weighing 40 ± 1 mg into centrifuge tubes, homogenizing with a steel ball in a ball mill, and centrifuging at 3000 rpm for 30 seconds at 4°C. After adding 400 μL of 80% methanol-water extraction agent and vortexing for 3 minutes, samples were subjected to freeze-thaw cycles in liquid nitrogen and dry ice, followed by vortexing and centrifugation at 12000 rpm for 10 minutes at 4°C. The supernatant was transferred and stored at -20°C before a final centrifugation. For HPLC analysis, samples were run on a Waters ACQUITY UPLC BEH C18 column at 40 °C with a flow rate of 0.4 mL/min, using a solvent system of water and acetonitrile with formic acid. The MS analysis was performed in IDA mode with specific source parameters and TOF MS scan settings. Data were processed using Proteo Wizard to convert to mzML format, XCMS for peak extraction and alignment, and R for PCA and HCA. Metabolites were identified through database searches and annotated with KEGG Compound and Pathway databases, with significant pathways determined by hypergeometric testing.

### In vitro proteomic analysis of biofilms

Protein extraction was performed via acetone precipitation, starting with grinding samples into powder and homogenizing in SDT buffer. After boiling, sonication, and centrifugation, protein solution was precipitated with acetone at -20°C. The precipitate was washed, dissolved in urea, and protein concentration was measured using a BCA kit. For digestion, proteins were reduced, alkylated, precipitated, and digested with trypsin overnight. Peptides were desalted and vacuum-dried before LC-MS/MS analysis on a nanoElute UHPLC system with a reverse-phase C18 column. The timsTOF Pro2 mass spectrometer operated in PASEF mode, acquiring MS and MS/MS spectra from 100 to 1700 m/z with a collision energy ramp. Raw data were analyzed with DIA-NN using a deep learning-based spectral library from the Homo sapiens SwissProt database, with an FDR of less than 1% for protein and ion identification, leading to quantification analysis.

## Results

### Metagenomics reveals microbiome profiles in pressure ulcers

#### Microbial composition

We collected swab samples from both pressure ulcer and normal skin patients (n=50) for 16S rRNA sequencing and metagenomic next-generation sequencing (mNGS). After removal of low-quality sequences and chimeras, we were left with a total of 3,113,803 high-quality sequences from all samples. Samples from group A (n=25) were collected from patients with pressure ulcers, whereas group B samples (n=25) were derived from normal skin (**Figure 1A**). We identified Proteobacteria, Actinobacteria, and Firmicutes as the top three phylum with the highest relative abundance (**Figure 1B**), while *Staphylococcus, Corynebacterium*, and *Klebsiella* were determined to be the most prevalent genera among the pressure ulcer group (**Figure 1C**).

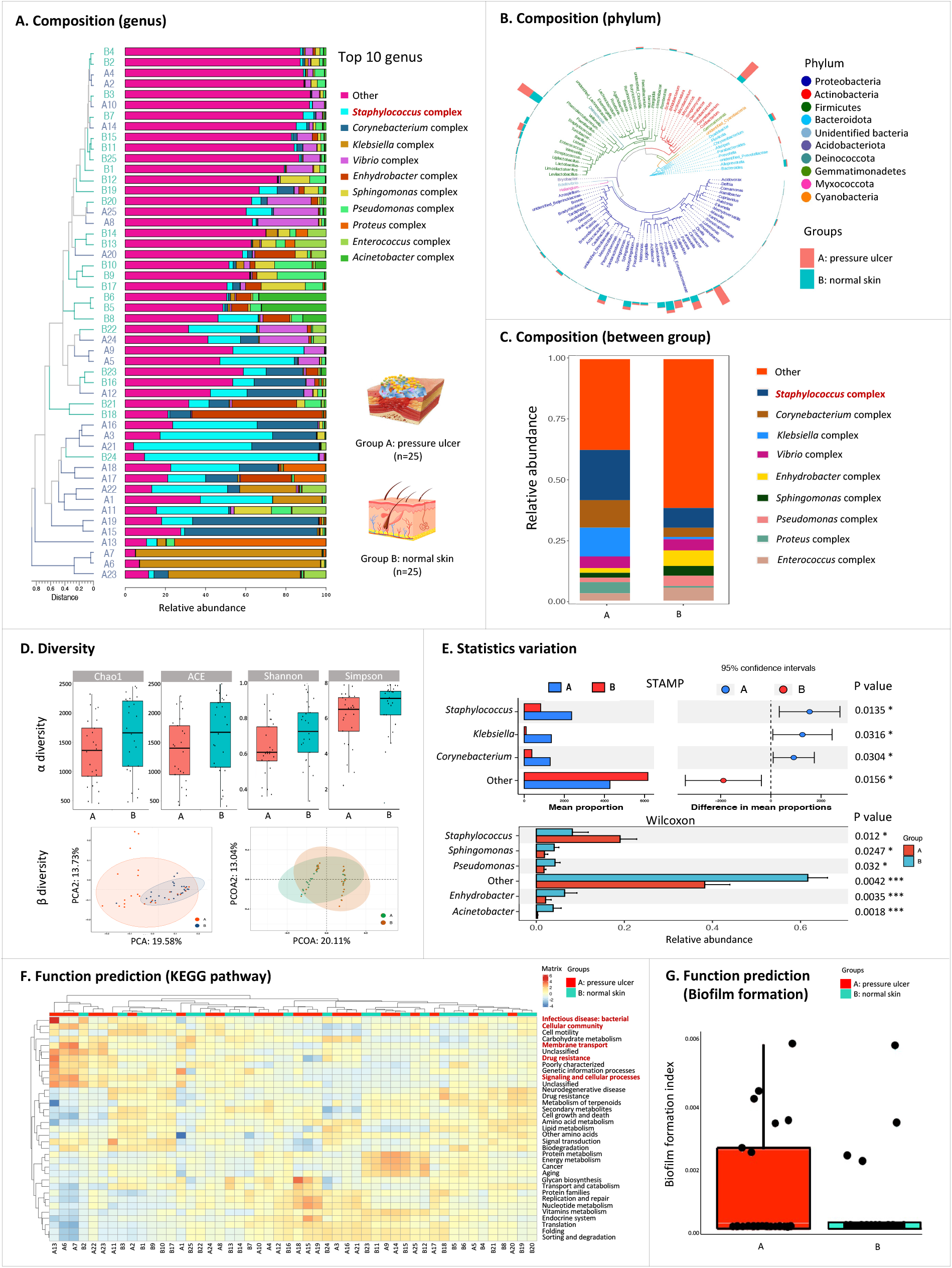

#### Diversity analysis

We proceeded to perform α-diversity analysis on the samples from both groups, indicating a substantial reduction in the richness and diversity of microbial communities within pressure ulcer as compared to normal skin. Concurrently, principal component analysis (PCA) and principal coordinate analysis (PCoA) demonstrated high reproducibility in the normal skin group, with samples forming distinct clusters that were separate from the majority of the pressure ulcer group samples (**Figure 1D**). Additionally, we utilized STAMP and Wilcoxon rank sum test to analyze the variance within microbial communities. The findings indicated that the most notable difference in microbial abundance was observed for *Staphylococcus* spp. (p=0.012∼0.0135), with significant disparities between the two groups, followed by *Klebsiella* (p=0.0316), *Corynebacterium* (p=0.0304) and others (**Figure 1E**).

#### Virulence factor prediction

The functional prediction analysis revealed that there were differences in pathogenic invasiveness and biofilm formation factors among the groups. In particular, samples from group A contained a set of functional genes associated with the induction of bacterial infectious diseases, cellular community, cell motility, drug resistance and signaling processes (**Figure 1F**). The Bugbase annotation results showed that the microbial community in group A may have increased biofilm formation capabilities, showing greater resistance to clinical antimicrobials compared to group B (**Figure 1G**).

#### Antimicrobial resistance prediction

A genus-level analysis of antimicrobial resistance genes (ARGs) between the pressure ulcer and normal skin groups revealed that *Staphylococcus* spp. strains in the samples possessed the highest number and diversity of ARGs (**Figure 2**). For instance, the aminoglycoside resistance gene *aadA*, macrolide resistance gene *ermB*, sulfonamide resistant gene *sul1*, tetracycline resistance gene *tetB*, as well as OXA beta-lactamases were present in higher quantities in the pressure ulcer group compared to the normal skin group. The aminoglycoside resistance gene *aac(6’)-1*, the β-lactam resistance gene *bacA*, and the macrolide resistance genes *ermX* showed peaks with high overlap and similar carriage across both groups. The presence of the *mecA* gene, linked to methicillin resistance, was identified in *Staphylococcus* species, while vancomycin resistance genes were found in *Enterococcus* species. This indicates that mixed wound infections involving bacteria with diverse antimicrobial resistance profiles can result in extensive resistance, especially to critical drugs such as vancomycin, which are considered the “last line of defense” in clinics [21].

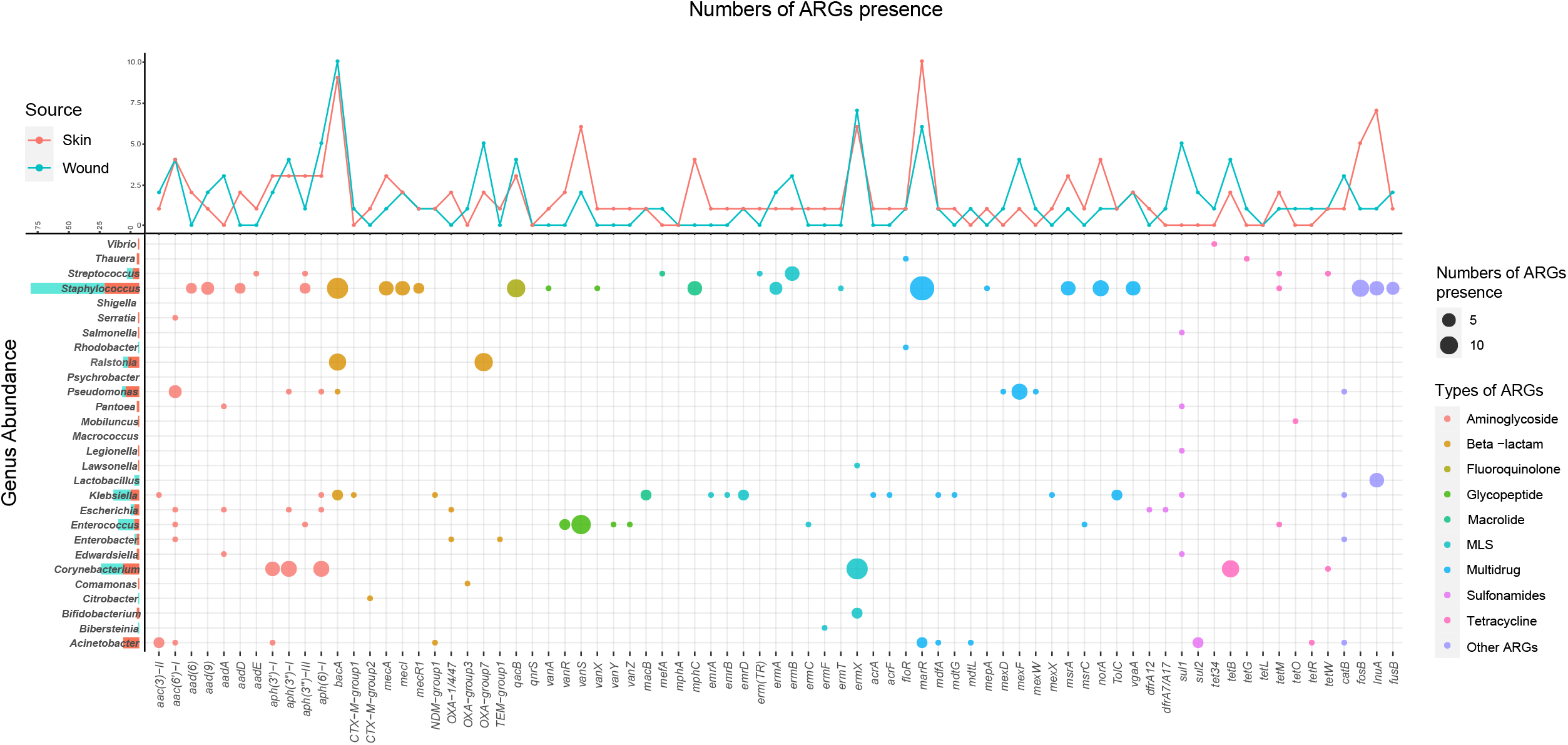

### Genomics exhibits characteristics of *Staphylococcus aureus* isolates

#### Bacterial whole genome analysis

To further character *S. aureus* present in pressure ulcers, we isolated 29 positive strains from pressure ulcer group for whole-genome sequencing (**Figure 3**). A total of 13 clonal complexes were identified among all *S. aureus* strains, which were further classified into 14 *spa* types and 13 ST types. The majority of the strains were affiliated with the CC22 clonal complex (n=8), followed by a proportion belonging to the CC59 clonal complex (n=4). The prediction of antimicrobial resistance genes and virulence factors revealed that the majority of *S. aureus* strains isolated from pressure ulcers possessed the *blaZ* (79.3%, 23/29) and *mecA* (48.3%, 14/29) genes, which are known to confer resistance to β-lactam antimicrobials. Furthermore, additional drug-resistant genes, including those for erythromycin, aminoglycoside, and tetracycline, were identified in these strains, highlighting the broad spectrum of antimicrobial resistance exhibited by *S. aureus* in wound infections. The prediction of virulence factors demonstrated that all *S. aureus* from pressure ulcer wounds contained the hemolysin encoding genes *hlgA, hlgB*, and *hlgC*. The *luk*F-PV and *luk*S-PV, which was the key virulence factors that cause tissue necrosis, were identified 48.3% strains (14/29), while human-associated *scn* gene was detected in 51.7% stains (15/29), highlighting the enhanced virulence of *S. aureus* isolated from pressure ulcer wounds.

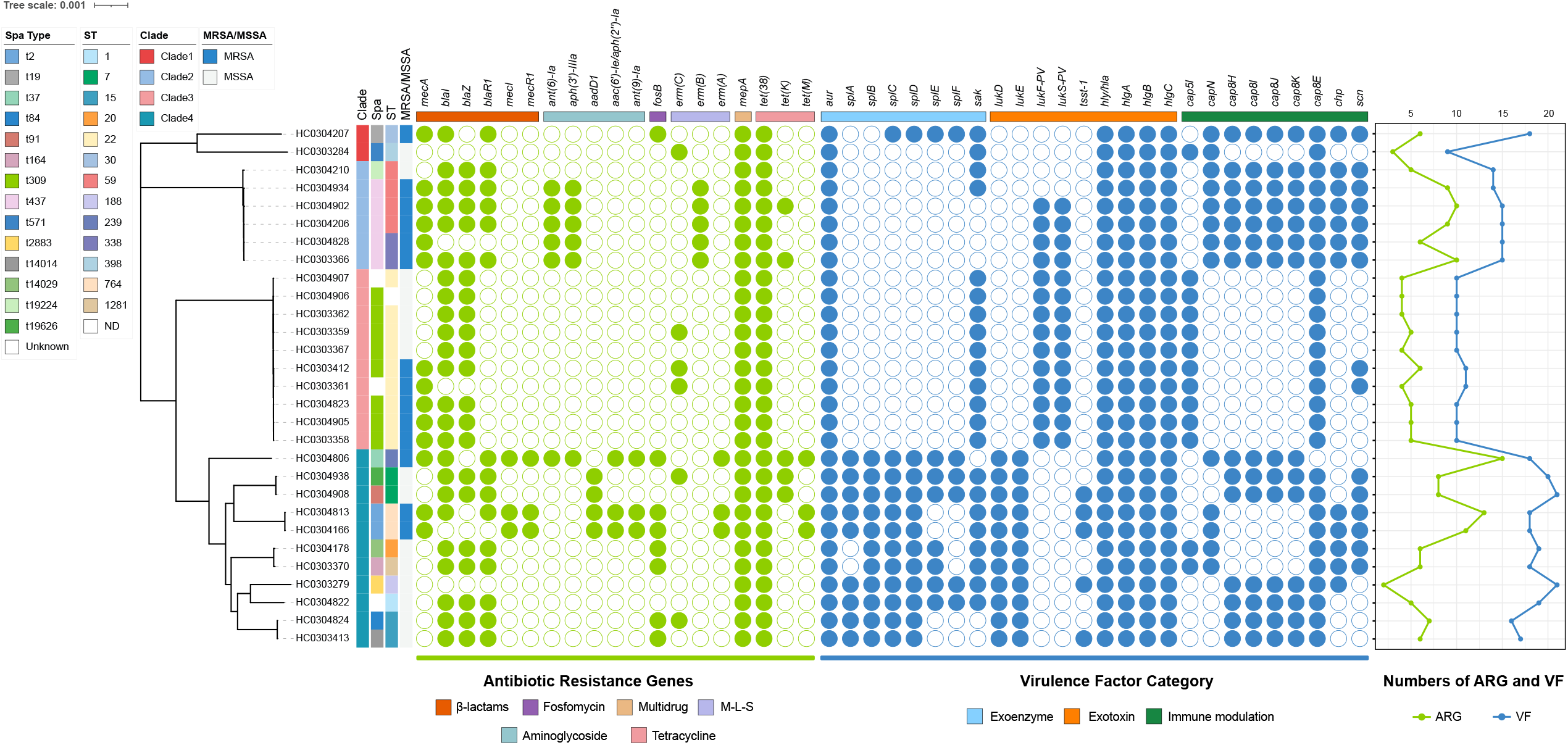

#### Skin-originated S. aureus lineages

We conducted a phylogenetic analysis on a total of 227 genomes, comprising 29 *S. aureus* isolates from this study and 198 skin-derived isolates sourced from public databases. The phylogenetic tree covers the period from 2003 to 2022 and includes isolates from a range of host environments (**Figure 4**). The 29 isolates from pressure ulcer wounds are distinctly marked in red on the phylogenetic tree. The analysis revealed that the majority of these isolates were distributed across different lineages, with one dominant cluster that affiliated with the CC22 clonal complex. Compared to *S. aureus* from other hosts, *fnbB* gene were only present in 13 strains isolated from pressure ulcers, nevertheless, *icaA icaB* and *icaD* genes were detected in 100%, 94.3%, and 99.6% strains. The varying presence and absence of biofilm genes among these isolates indicate the complexity of microbiome infections. The effects of other bacteria with strong biofilm-forming ability should also be considered in addition to *S. aureus*.

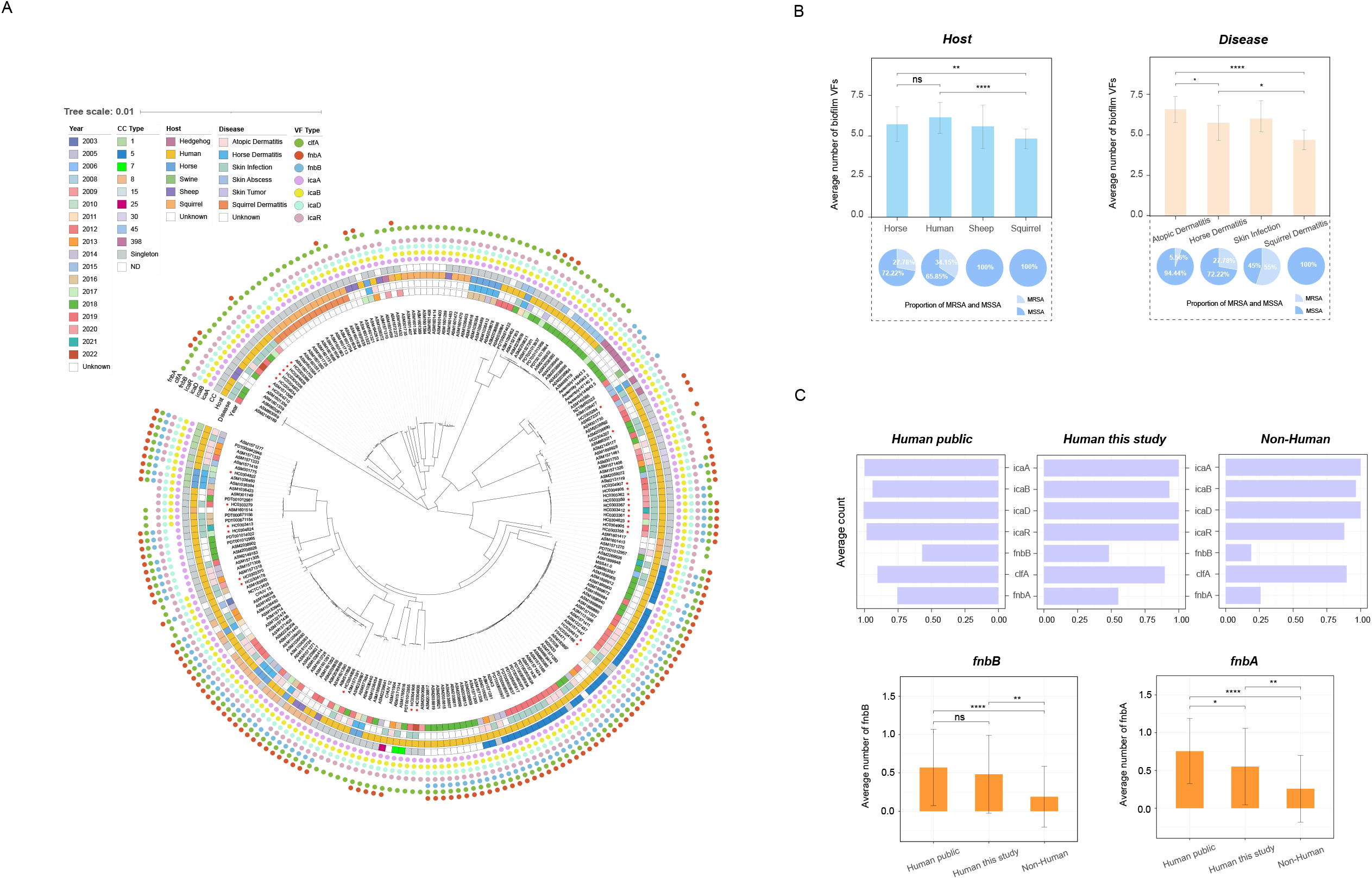

### Histopathology observes *Staphylococcus aureus* in biofilm structure

#### Section staining analysis

The combined results from Hematoxylin-Eosin, Masson, and Sirius-red staining indicated the presence of localized accumulations of collagen fibers and proteins, which are essential elements in the formation of biofilms in pressure ulcers group (**Figure 5A/B/C**). Additionally, gram staining results revealed a dense clustering of gram-positive bacteria and PAS staining showed an abundance of polysaccharides in the area, which, along with extracellular nucleic acids, constitute a significant part of the biofilm structure (**Figure 5D/E**).

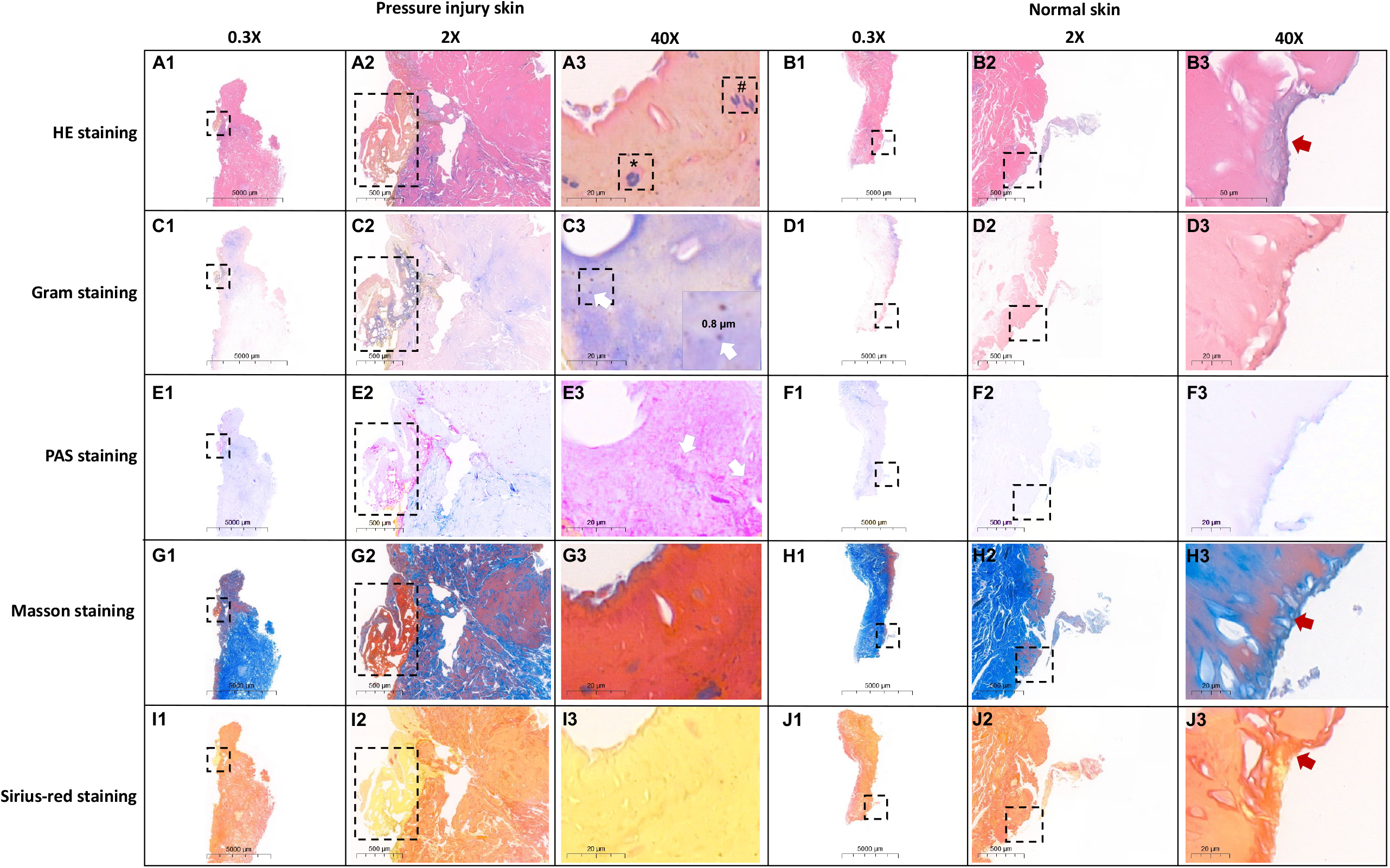

#### Fluorescence in situ hybridization

To elucidate the presence and spatial distribution of the microbial communities within the biofilm structure of pressure ulcer wounds, we employed the eub388 probe, which targets the *S. aureus agrBDCA* operon, to conduct FISH on samples from the pressure ulcer group. DAPI and conserved region of 16S rRNA were employed to show the general cells and bacteria. The FISH signal detection corroborated the section staining results, confirming the presence of a significant number of *S. aureus* in the biofilms diffusely distributed and formed to be clustered as a “strip or rotund-shaped” structure of strongly positive signals attached cells (**Figure 5F**). This clustering is hypothesized to potentially enhance the adaptability of the microbial community and the stability of the biofilm.

### Combined metabolomics and proteomics explore the enriched pathways in biofilm formation

#### In vitro biofilm formation test

To further understand the interaction between *S. aureus* isolated from pressure ulcer wounds and its biofilm, we conducted an in vitro biofilm formation experiment. Statistical analysis showed that these 29 strains of showed various degrees of biofilm positive or weak positive formation (**Figure 6A/B**). Among them, strain numbers 571, 725 and 892 were identified as biofilm positive according to OD595 values (OD=2.28, 2.46, 2.81), and strain numbers 812, 817 and 821 were determined as weak biofilm signal (OD=0.21, 0.50, 0.24), with standard strain ATCC25923 as control (OD=0.30, **Figure 6C**).

#### Metabolomics and proteomic analysis

Our proteomics and metabolomics analysis revealed significant insights into the interactions between *S. aureus* strains from pressure ulcer wounds and their biofilm formation. The PCA analysis revealed that the two high biofilm-producing strains, A2 and A3, formed a cluster, while A1 and three low biofilm-producing strains from group B were grouped together. The ATCC standard strains showed distinct differences from the strains in groups A and B in terms of their proteomic and metabolomic profiles (**Figure 6D**).

The proteins that differed between samples in group A and group B were primarily located in the cytoplasm and cytoplasmic membrane (**Figure 6E**). The GO annotation results for proteins revealed that a significant portion of the differential proteins between samples from group A and group B were associated with metabolic processes, catalytic functions, and membrane composition. The differential combination results among groups A, B, and C were similar to the comparison results between A and B. However, the number of annotated differential proteins was larger, particularly the number of membrane-related differential proteins in the comparison between group C and A was higher than that between group C and B (**Figure 6F**). It is hypothesized that this may be related to the fact that the biofilm strength of the strains in group A is higher than that of the strains in group B.

Furthermore, based on the KEGG enrichment analysis of differential metabolites and proteins, we identified KEGG pathways that were commonly enriched in both proteomics and metabolomics for the sample combinations of groups A and B (**Figure 6G**). The results indicated that numerous metabolites were significantly enriched in metabolic pathways, including amino acid biosynthesis and secondary metabolite biosynthesis. Additionally, differential proteins were significantly enriched in pathways such as ABC transporter proteins, purine metabolism, and nucleotide metabolism. The ABC family multidrug transporter proteins facilitated the efflux of various cytotoxic drugs across the cell membrane [22], making them a key factor in the development of drug resistance in *S. aureus*. Apart from ABC transporter proteins, all the enriched pathways mentioned above were directly or indirectly involved in the formation of *S. aureus* biofilms, which significantly enhanced the bacterial ability to adapt to adverse environments and increased their survival rate under challenging conditions.

## Discussion

The delayed healing of chronic wounds, such as pressure ulcers, is a complex phenomenon influenced by numerous factors including microbial composition and biofilm formation. Previous study has identified *S. aureus* as a primary pathogen in wound infections [23], nevertheless, there is a lack of data on the colonization of *S. aureus* in chronic wounds. To address this knowledge gap, we used a multi-omics approach that included genomic, proteomic, and metabolomic analyses to elucidate the microbial composition and in-depth characterization of *Staphylococcus aureus* isolated from pressure ulcers.

Chronic wounds, as opposed to acute ones, are significantly more prone to biofilm formation (>60%). They also exhibit a notable reduction in microbial diversity, characterized by a predominance of pathogenic bacteria such as *Staphylococcus, Pseudomonas, Corynebacterium, Streptococcus*, and *Enterococcus* [24-27]. In our research, *Staphylococcus* emerged as the most common species detected, trailed by *Klebsiella, Proteus*, and *Corynebacterium*, aligning with previous research observations.

In addtion, the identified *Staphylococcus, Corynebacterium, Klebsiella*, and *Enterococcus* in this study harbored a significant number of antimicrobial resistance genes, with a particular prevalence of the *bacA* gene in *Staphylococcus* and *Klebsiella*. The *bacA* gene encodes an inner membrane protein that plays a role in sustaining chronic infections across various host-pathogen interactions [28], with the prevalence from 10% to 59% in chronic wounds. Our findings highlight the importance of antimicrobial management and testing for resistance in patients with pressure ulcers.

*Staphylococcus aureus* frequently develops biofilms within chronic wounds, leading to increased antimicrobial resistance [29,30]. Our study utilized whole genome sequencing to analyze 29 *S. aureus* isolates and discovered that approximately 79% of these strains possessed *blaZ*, which is responsible for the production of penicillinase [31]. Furthermore, 50% of the strains exhibited resistance to methicillin, which were classified as methicillin-resistant *Staphylococcus aureus* (MRSA) strains. Furthermore, the presence of the *scn* gene in these isolates suggests that the infections may originate from human-to-human transmission, especially for elderly patients who have been bedridden in hospitals over long periods.

According to the fluorescence in situ hybridization, it can be confirmed that *S. aureus* biofilms were formed in the wounds of patients with pressure ulcers in this study, which is consistent with previous studies. Interestingly, Fazli et al. found that bacteria in chronic wounds tend to aggregate and form dense clusters [32]. A similar phenomenon was also found in our study that *S. aureus* was mostly distributed in clusters and colonized in areas close to the surface of chronic wounds. However, further research was needed to illustrate the bacterial colonization and distribution deep within wounds.

*Staphylococcus aureus* CC22 is a lineage that predominantly affects humans and is recognized as one of the top five MRSA clone complexes reported globally, frequently implicated in hospital-acquired infections [33-35]. Among the *S. aureus* collected from the patients with pressure ulcers in this study, CC22 accounted for the highest proportion and clustered together on the evolutionary tree, indicating a possibility of hospital infection or a small-scale outbreak. Compared to other virulence factors, pvl is only present in a tiny proportion of strains, according to earlier research. However, all the CC22 strains in our investigation were able to secrete leukocidin, indicating that highly pathogenic *S. aureus* is linked to the microbial population of pressure ulcer. Additionally, the CC59 strains in this study are the second most prevalent. It has been documented that CC59-ST59 is a significant lineage associated with the spread of methicillin resistance among Staphylococcus aureus in Chinese clinical settings [36,37].

Finally, we performed an experiment on the biofilm formation of isolated *S. aureus* strains and integrated it with a comprehensive analysis of proteomics and metabolomics data. Our study, based on the biofilm formation experiment, conducted a comparative analysis between *S. aureus* strains with different biofilm formation ability. The results showed that differential proteins, metabolites, and enriched pathways are closely related to the formation of biofilms to benefit bacterial adaptation to improve survival rate in the hospital selective settings.

Limitations of this work are that a larger cohort could enhance the statistical power and applicability of our findings. Secondly, our study primarily focused on the presence and characteristics of S. aureus, which may not capture the full spectrum of microbial interactions within the wound environment. Future studies could benefit from a more comprehensive analysis of the broader microbial community.

In conclusion, our study has unveiled the overall profile of microorganisms colonizing pressure ulcer wounds, encompassing an analysis of bacterial composition and functional genes, and has highlighted notable disparities between the pressure ulcer and the control group. The substantial presence of *S. aureus* within the biofilm was validated using pathological tissue sections and FISH techniques. We have deciphered the pathways and metabolites of *S. aureus* in pressure ulcers via proteomics and metabolomics approach bioinformatically. Our research offers a scientific foundation for understanding the colonization patterns of *S. aureus* biofilms in pressure ulcers wounds.

## Author contributions

CG, GJ, YL, JZ, JL, SL performed the analysis. CG, HL, ZZ wrote the main manuscript text. XW, ZZ, EX, HL prepared the data. All authors reviewed the manuscript.

## Institutional review board statement

The experiment was strictly conducted according to the Guide for Care and Use from the Research Ethics Committee of Soochow University (20210220). All procedures involving human participants were performed to the ethical standards.

## Informed consent statement

Informed consent was obtained from all subjects involved in the study.

## Data availability statement

The metagenomic dataset and 16S rDNA dataset in this study have been deposited in the CNCB database with GSA accession numbers CRA021958 and CRA022283, respectively. The Staphylococcus aureus dataset has been deposited in the NCBI database with BioProject accession number PRJNA1190906.

